# Linking dendroecology and association genetics: Stress responses archived in tree rings associate with SNP genotypes in *Abies alba (Mill.)*

**DOI:** 10.1101/125302

**Authors:** Katrin Heer, David Behringer, Alma Piermattei, Claus Bässler, Bruno Fady, Hans Jehl, Sascha Liepelt, Sven Lorch, Andrea Piotti, Giovanni Guiseppe Vendramin, Max Weller, Birgit Ziegenhagen, Ulf Büntgen, Lars Opgenoorth

**Affiliations:** Philipps-University Marburg, Faculty of Biology, Conservation Biology, Marburg, Germany; Marche Polytechnic University, Department of Agricultural, Food and Environmental Sciences, Ancona, Italy; DendroScience, Swiss Federal Research Institute WSL, Zürcherstrasse 111, 8903 Birmensdorf, Switzerland; Bavarian Forest National Park, Freyunger Str. 2, Grafenau, Germany; INRA, UR Ecologie des Forêts Méditerranéennes, Avignon, France; Philipps-University Marburg, Faculty of Biology, Department of Ecology, Marburg, Germany; National Research Council, Institute of Biosciences and Bioresources, Sesto Fiorentino, Firenze, Italy; Department of Geography, University of Cambridge, Downing Place, CB2 3EN Cambridge, UK; CzechGlobe, Global Change Research Institute CAS and Masaryk University, Kotlářská 2, 61137 Brno, Czech Republic

**Keywords:** Candidate genes, dendrophenotypes, genetic association, random forest, silver fir, SO_2_ pollution

## Abstract

- Genetic association studies in forest tress would greatly benefit from information on tree response to environmental stressors over time. Dendroecology can close this gap by providing such time series measurements. Here, we jointly analyzed dendroecological and genetic data to explore the genetic basis of resistance, recovery and resilience to episodic stress in silver fir.
- We used individual level tree-ring data to characterize the growth patterns of surviving silver fir (*Abies alba*) during the forest dieback in the 1970s and 1980s in Central Europe and associated them with SNPs in candidate genes.
- Most trees at our study sites in the Bavarian Forest experienced severe growth decline from 1974 until the mid-1980s, which peaked during the drought year of 1976. Using the machine learning algorithm random forest, we identified 15 candidate genes that were associated with the variance in resistance, resilience and recovery among trees in this period.
- With our study we show that the unique possibility of phenotypic time series archived in tree-rings are a powerful resource in genetic association studies. We call for a closer collaboration of dendroceologists and forest geneticists to focus on integrating individual tree level signals in genetic association studies in long lived trees.

## Introduction

Genetic association studies in forest trees often focus on phenotypes related to stress events (e.g. González-Martínez *et al*., 2006; Budde *et al*., 2014), with the goal of understanding the genetic basis of stress responses in the light of global change. Episodic stress events, such as droughts and storms, are of particular interest as they are expected to significantly increase during the 21^st^ century due to human-induced global climate change (IPCC, 2014). However, recording plant responses to episodic stresses in natural conditions is notoriously difficult due to the unforeseeable timing of such events. Here we utilize the fact that trees archive their reaction to environmental conditions in their wood anatomical structure and annual growth rings (Fritts & Swetnam, 1989). Phenotypic measures derived from wood cores include e.g. ring width, anatomical features, isotopic ratios or wood density, all of them available in dated time-series. We call all such traits derived from dated annual growth rings that are available as time series “dendrophenotypes” and propose that they can be a prime resource in genetic association studies. However, since classical dendrochronological studies usually discard the variation of dendrophenotypic traits among individuals as noise (Buras *et al*., 2016), a shift of focus towards this individual level variation is needed. In this study we present for the first time the association of SNP genotypes from stress related candidate genes with dendrophenotypes.

A prime example of an episodic stress event in Europe was marked by the forest dieback in the 1970s and 1980s (Kandler & Innes, 1995), which has been widely attributed to air pollution, particularly SO_2_ and O_3_ emissions, in combination with a series of exceptionally dry years (Elling *et al*., 2009). The conifer *Abies alba* Mill. (silver fir) was severely affected by these conditions with large scale dieback and severe growth decline (Kandler & Innes, 1995). This massive stress episode provides an ideal test case to trace intraspecific variation in dendrophenotypic measures and associate them with candidate genes on small spatial scales in a natural population. For this purpose we characterized the growth decline of *A. alba* in the 1970s and 1980s by applying the resilience concept of Lloret *et al*. (2011). To do so we obtained dendrophenotypes from wood cores for 193 *A. alba* trees from the Bavarian Forest National Park and associated them with SNPs at candidate genes mainly related to stress reactions (Roschanski *et al*., 2016). Since growth as a quantitative trait is likely influenced by many genes we not only applied a single-locus approach for the genetic association, but also used a random forest analysis to capture both the marginal effect of single SNPs, as well as the combined effect of multiple SNPs on a phenotype.

Since the stress episode caused significant dieback we expect that a large majority of the surviving trees exhibit a marked growth decline in the 1970s and 1980s, but with a high level of variance among individuals regarding resistance, resilience and recovery. Further, we expect the variation in the dendrophenotypes to be associated with genetic variation in a number of the targeted genes, and thus, to confirm them as candidate genes for stress response. Finally, we discuss the potential of dendrophenotypes in genetic association studies for exploring the genetic basis of growth decline and resilience in episodic stress scenarios in the context of climate change.

## Material and Methods

### Study site

*Abies alba* trees were sampled and monitored in two sites in the Bavarian Forest National Park, Germany which are situated at opposite ends of an elevational gradient but within gene flow distance to maximize phenotypic variation and minimize drift effects at the same time (Lotterhos & Whitlock, 2015). Our sampling sites were located at 770 m a.s.l. (*Filzwald;* 48.929°N, 13.406°E) and 1.120 m a.s.l (*Rachelsee;* 48.975°N, 13.400°E) on the southern slope of Mt. Rachel (Fig. 1). Mean annual temperature varies between 3.8°C and 5.8°C with a mean annual precipitation from 1,200 to 1,800 mm in the National Park (Bässler, 2004). Here, silver fir grows in mixed mountain forests in combination with *Fagus sylvatica* and *Picea abies* which is the natural vegetation at elevations below 1,150 m. At each site, 100 adult silver fir trees were georeferenced and permanently marked with numbered tags. Temperature and humidity were recorded at both sampling sites with data loggers (DK320 DM HumiLog, Drieβen & Kern, Bad Bramstedt, Germany) starting in spring 2014. The lower elevation sampling site (‘low site’ hereafter) is characterized by flat terrain and subjected to accumulating cold air from higher elevations, which leads to frequent late and early frost events in spring and autumn, respectively. In contrast, the sampling site at higher elevation (‘high site’ hereafter) is located on a steep slope surrounding the Rachel lake. The lake influences the local climatic conditions by buffering cold temperatures. Therefore, early fall and late spring frost events are less frequent and maximum temperatures are lower than at the low site. Mean temperature, however, did not significantly differ between sites during our study period (Table S1).

**Figure 1.**
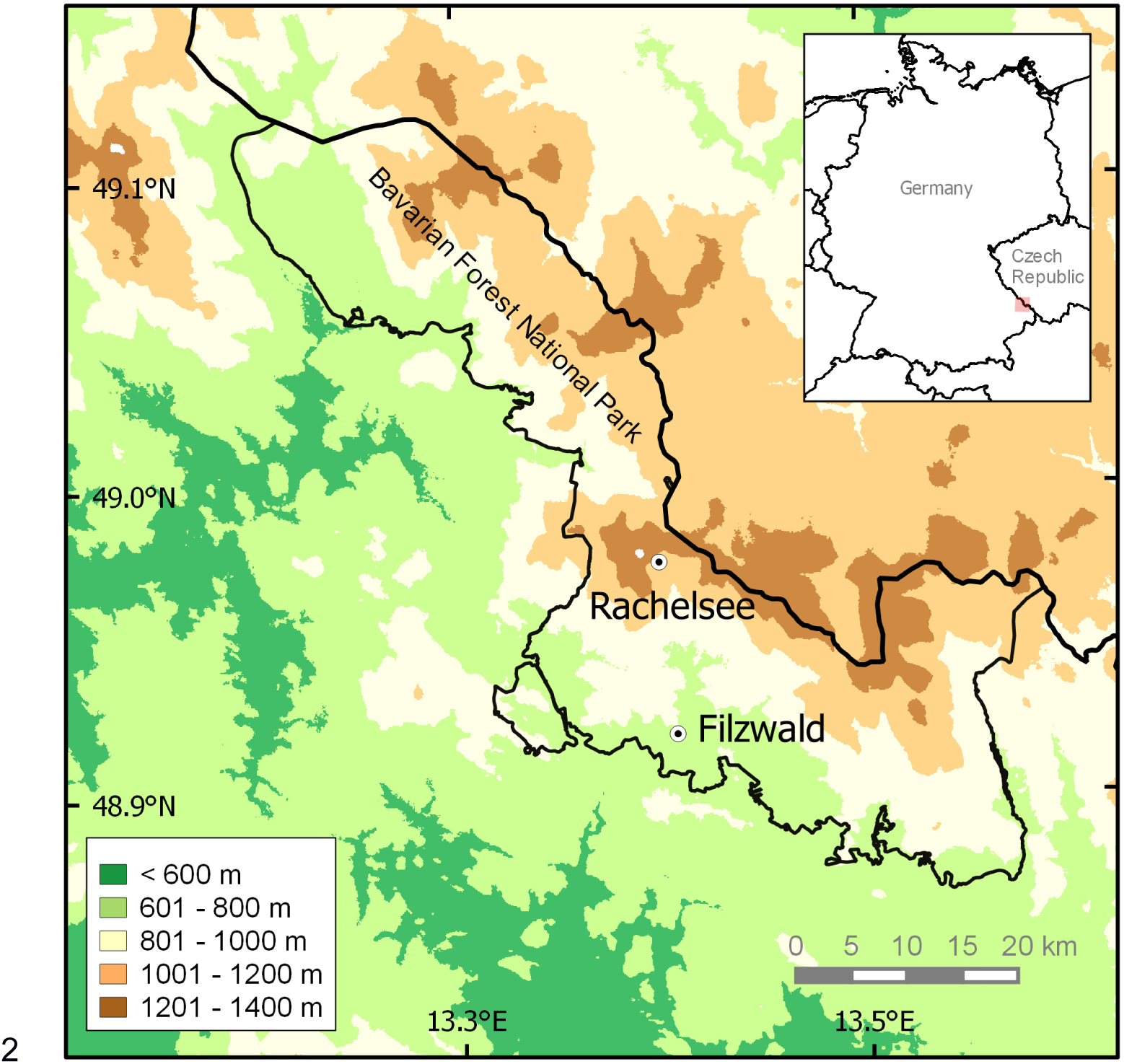
Study site in the Bavarian Forest National Park with the sampling sites Rachelsee (‘high site’, 1120 m a.s.l.) and Filzwald (‘low site’, 770 m a.s.l.)

### Phenotypes from wood cores

To obtain data on tree-ring width (TRW), we extracted two wood cores per tree at breast height with an increment borer. If trees grew on slopes, they were cored at a 90° angle to the slope to avoid compression wood. After drying, intact or slightly fractured wood cores were cut with a microtome (WSL, Birmensdorf, Switzerland) to obtain a smooth surface. The contrast between earlywood and latewood was enhanced with chalk. Cores that had several fractures (14 out of 375) were mounted on wooden holders and smoothed with sandpaper. TRW was measured with a precision of 0.01 mm using a LINTAB digital positioning table whose movements were transmitted to the TSAP-Win Scientific Software (Rinntech, Heidelberg). A master series for each site using COFECHA (Grissino-Mayer, 2001) was constructed and each series was cross-dated against this master series to avoid dating errors due to missing rings. We obtained reliable data from a total of 375 cores from 193 trees.

All tree ring time series were standardized to a mean value of one to obtain a dimensionless tree-ring index (TRI) using the *detrend* function in the R package dplR (Bunn, 2008; R Core Team, 2016). Based on the inspection of chronologies at the site level, we identified the years from 1974 to 1983 as the period where most trees exhibited the strongest growth decline. In the following, we refer to this period as “depression period” and to the ten years before as “reference period” (Fig. 3). For the dendrophenotypic characterization of individual trees, we determined their resistance, recovery and resilience (following Lloret *et al*., 2011) to possible effects of airborne SO_2_ pollution (Fig. 2). In this framework, resistance describes the ratio of TRI during vs. before the extreme event; recovery describes the ratio of TRI after vs. during the event; and resilience describes the ratio of TRI after vs. before the event. Based on these definitions, we calculated the following dendrophenotypic traits: (1) the steepness of the start of the depression period in 1974 (*slope_73_74*) was defined as the slope of the standardized TRI between the years 1973 and 1974; (2) the resistance in the depression period (*resistance_dep*) was defined as the ratio between the average TRI from 1964 to 1973 and the average TRI during the depression period (1974-1983); (3) the recovery after the depression period (*recovery_dep*) was calculated as the ratio between the average TRI in the ten years after 1983 and the average TRI during the depression period, and (4) the end of the individual growth depression (*end_dep*) was defined as the year when growth surpassed the level during the reference period. To calculate the latter, we compared mean TRI after 1973 in a moving window of three years to the mean growth in the reference period (Fig. 2) and determined the year when the mean of the moving window first surpassed the mean of the reference period.

**Figure 2.**
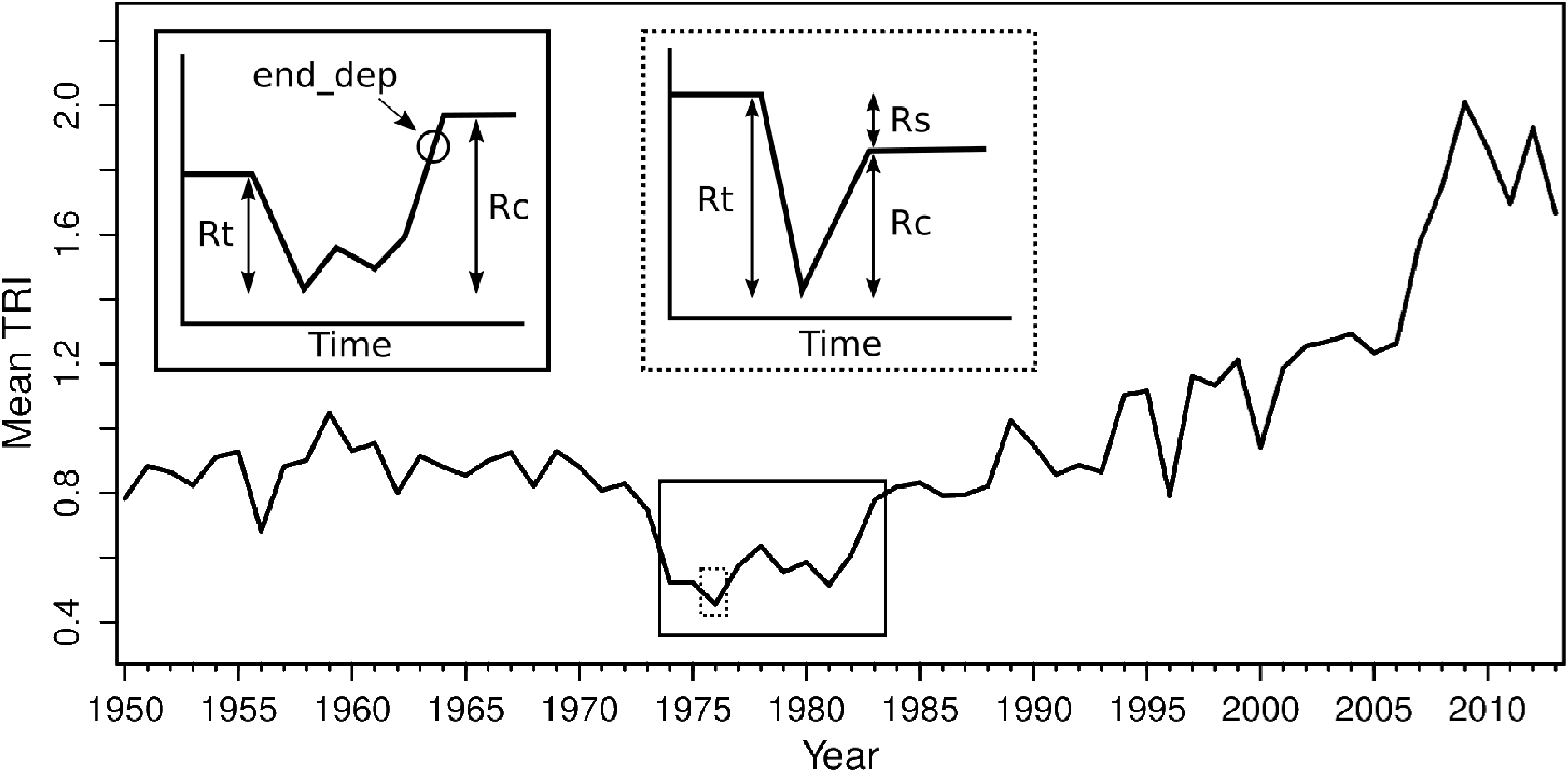
Mean tree-ring index (TRI) of all individuals across both sites for each year in the period from 1950 to 2013. The insets (modified after Lloret et al. 2011) are simplified graphical representations of the dendropheno-typic measures resistance (Rt), recovery (Rc), resilience (Rs) and end of depression period (end_dep) for the growth depression (solid box) and for the drought year 1976 (dashed box). For details regarding the calculation of the indices see material and methods.

**Figure 3.**
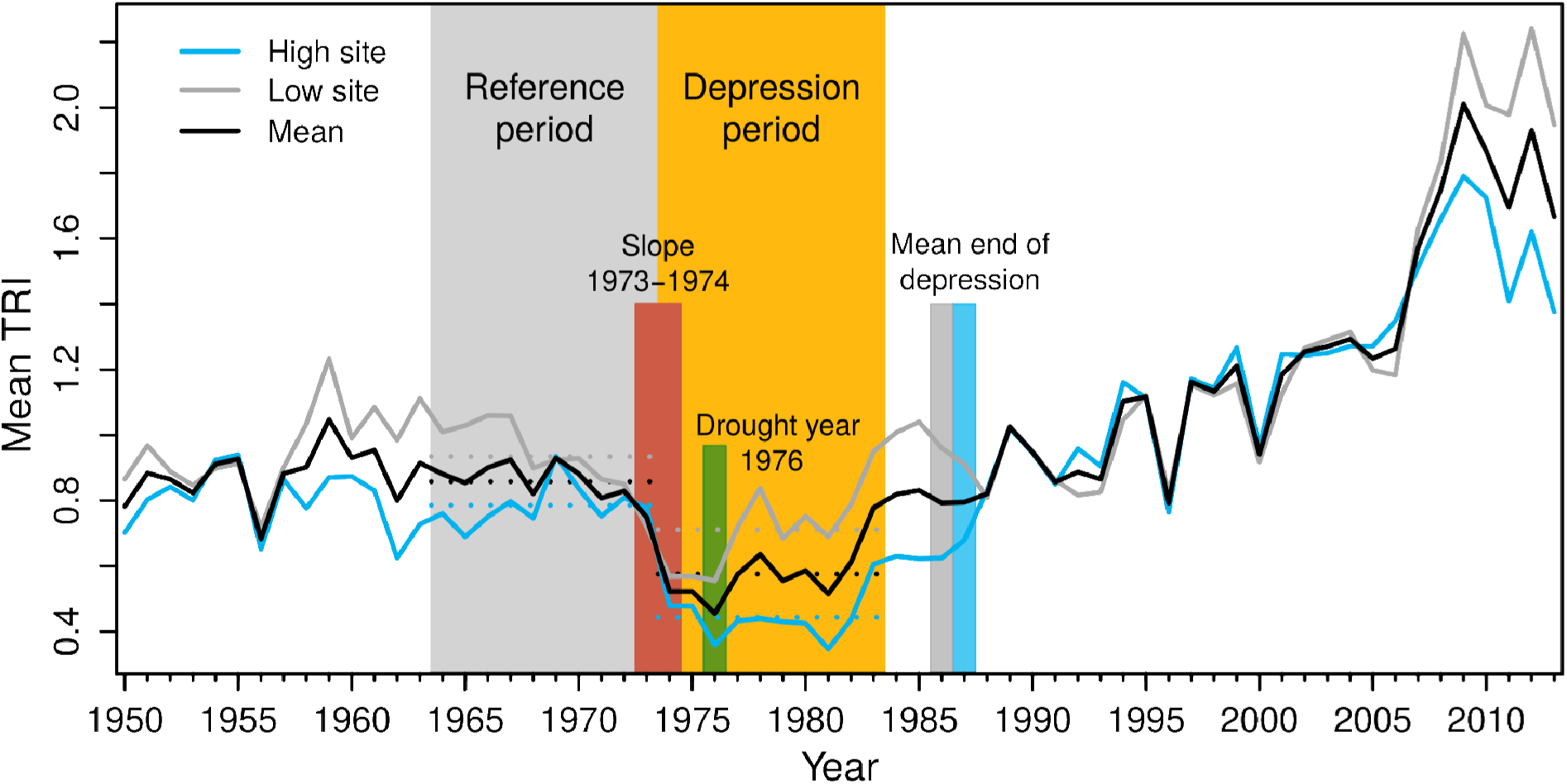
Mean tree-ring index (TRI) of all individuals across both sites and separated for the two sites (high and low) for each year in the period from 1950 to 2013.The shaded reference and depression period are the basis for the calculation of the resistance to SO_2_ pollution. Dotted lines represent the mean TRI for the corresponding period and site. Vertical bars from left to right mark the onset of the depression period for which we calculated the slope, the drought year 1976, and the mean end of the growth depression for low and high elevations, respectively.

In addition, we focused on the individual growth reaction in the year 1976, which has been identified as one of the driest summers in Europe (Briffa *et al*., 2009). We used the *res.comp* function in the R package *pointRes* (van der Maaten-Theunissen *et al*., 2015) to calculate (5) the *recovery_1976*, (6) *resilience_1976* and (7) *resistance_1976* of each tree towards the drought conditions in 1976. For the calculation, we considered a two-year window to take into account the already reduced growth in the two years prior to 1976 within the period of growth depression (Lloret *et al*., 2011). To compare dendrophenotypes between sites, we calculated mean values and corresponding standard deviations (SD) for each site and used Welch’s unequal variances t-tests to compare them. For the genetic association, all dendrophenotypes were centered and scaled within each site to exclude confounding effects due to environmental and site conditions, using the *scale* function in the R package base with default parameters.

### Genotyping

For DNA extraction, fresh needles were collected from each tree, and immediately dried on silica gel. Needles were sent to LGC Genomics (Middlesex, United Kingdom) for genotyping. Using KASP assays, we targeted 267 polymorphic and functionally annotated SNPs in candidate genes whose selection is described in detail by Roschanski *et al*. (2013, 2016). Briefly, genes were selected based on a literature search that included the keywords adaptation, candidates, drought evolution RT-PCR and selection, or based on drought-related Gene Ontology terms. Out of these 267 SNPs, 241 could be successfully genotyped using the KASP technology in our samples.

The dataset was initially filtered by removing SNPs that were not called correctly for the majority of the data set (> 80 % missing data) because they could interfere with subsequent filtering steps. Subsequently, individuals and SNPs with more than 10 % missing data, as well as monomorphic SNPs, were removed. We then selected all SNPs with a minor allele frequency > 3 %. All SNPs were tested for pairwise linkage disequilibrium (LD) using the Genome Variation Server 147 v. 12.00 (National Heart, Lung, and Blood Institute, http://gvs.gs.washington.edu/GVS147/index.jsp). If pairs of SNPs were tightly linked (r^2^ = 1) one of the SNPs was removed. This was only the case for SNPs located on the same contig of the transcriptome assembly (see Roschanski *et al*., 2013).

With this set of SNPs, we imputed missing genotypes using Beagle 4.1 (Browning & Browning, 2016) without using a reference sequence. After imputation, the dataset was cleaned again using the same filtering steps as described above. Finally, all SNPs with less than five individuals per allele combination were removed from the dataset to ensure sufficient replication, resulting in 193 individuals and 130 SNPs from 103 genes.

### Population clustering

As the methods we used for phenotype-genotype association are sensitive to population structure, we applied two approaches to determine whether we could detect a genetic structure within or between sampling sites. First, we used the Bayesian clustering algorithm implemented in STRUCTURE 2.3.4 (Pritchard *et al*., 2000) based on the admixture model with correlated allele frequencies. We set the burn-in to 10^5^ iterations followed by 5 × 10^5^ MCMC repetitions. We conducted 10 runs for each K from 1 to 6. Second, we conducted a PCA as implemented in the R package *adegenet* (Jombart & Ahmed, 2011).

### Genetic association analysis

For the association analysis of SNPs and dendrophenotypes, two approaches were used. First, we applied a frequently used univariate approach, namely general linear models (GLMs) as implemented in TASSEL v. 5.0 (Bradbury *et al*., 2007), with each SNP as the independent variable and each dendrophenotype as the response variable. For each dendrophenotype, we ran GLMs with 10,000 permutations to obtain p-values independent of the data distribution. The threshold for statistical significance at 5% was 0.007 after Bonferroni multiple test correction (0.05/7).

Univariate approaches only take into account a single SNP per test. However, most phenotypes are influenced by multiple genes with small effects. Thus, we also applied the machine learning algorithm “random forest” which captures both marginal and interaction effects among SNPs. In this study, we used the feature selection procedures as implemented in the R packages Boruta (Kursa *et al*., 2010) and VSURF (Genuer *et al*., 2015). A detailed description of these methods is provided in the supplement (Methods S1).

Although SNPs were already annotated for a previous publication (Roschanski *et al*., 2016), we repeated this step, as the information in the NCBI database is constantly updated. All SNPs that could be associated with the dendrophenotypes with more than one of the above mentioned methods were compared to known sequences from NCBI’s GenBank nonredundant protein database (NR) using the translated BLAST algorithm (blastx v. 2.6.1+, Altschul *et al*., 1997). The best hit, based on the Expect value that provided a functional annotation, was selected for each gene and the corresponding biological process keywords were retrieved from the Gene Ontology (GO) database (UniProtKB, The UniProt Consortium, 2015).

## Results

### Dendrophenotypes

We obtained data on dendrophenotypes for 98 and 95 individuals from the high and low site, respectively. Almost all trees from the high plot (97%) were affected by a growth decline, whereas only 80% of the low site individuals showed such a reaction (Fig. 4B). For *end_dep* we determined mean values of 1986 and 1987 for the low and high site, respectively (Fig. 4D). Individuals from the high site had significantly lower values for *slope_73_74* and *resistance_dep*, and on average, significantly higher values for *recovery_dep* when compared to the low site (Fig. 2, Fig. 4, Table 1). Most trees the high site showed a marked growth decline in 1976, while trees from the low site showed, on average, no further decline which resulted in a significantly higher value of *resistance_1976* (Fig. 4F). At the low site, trees already showed an increase in growth after 1976 which is reflected in the higher values for *recovery_1976* and *resilience_1976*, although only the latter differed significantly between sites (Fig 4E,G).

**Table 1.** Summary statistics for the comparison of dendrophenotypes between the two sites. Test results from a Welch’s *t*-tests are provided.

| Site | *slope_73_74* | *resistance_dep* | *recovery_dep* | *end_dep* | *resilience_1976* | *resistance_1976* | *recovery_1976* |
| --- | --- | --- | --- | --- | --- | --- | --- |
| Low (mean ± SD) | -0.15 ± 0.13 | 0.82 ± 0.67 | 1.66 ± 1.11 | 1986.49 ± 11.21 | 1.37 ± 0.41 | 0.97 ± 0.30 | 1.59 ± 1.76 |
| High (mean ± SD) | -0.30 ± 0.20 | 0.58 ± 0.21 | 1.96 ± 0.89 | 1987.48 ± 6.57 | 0.95 ± 0.30 | 0.76 ± 0.22 | 1.41 ± 1.37 |
| *T* | -5.983 | -3.3264 | 2.0079 | 0.72087 | -7.9174 | -5.4393 | -0.81124 |
| Df | 169 | 109 | 179 | 146 | 171 | 173 | 176 |
| *p*-value | **<0.001** | **0.0012** | **0.046** | 0.47 | **<0.001** | **<0.001** | 0.48 |

SD: standard deviation, *t*: *t*-statistic, df: degrees of freedom

**Figure 4.**
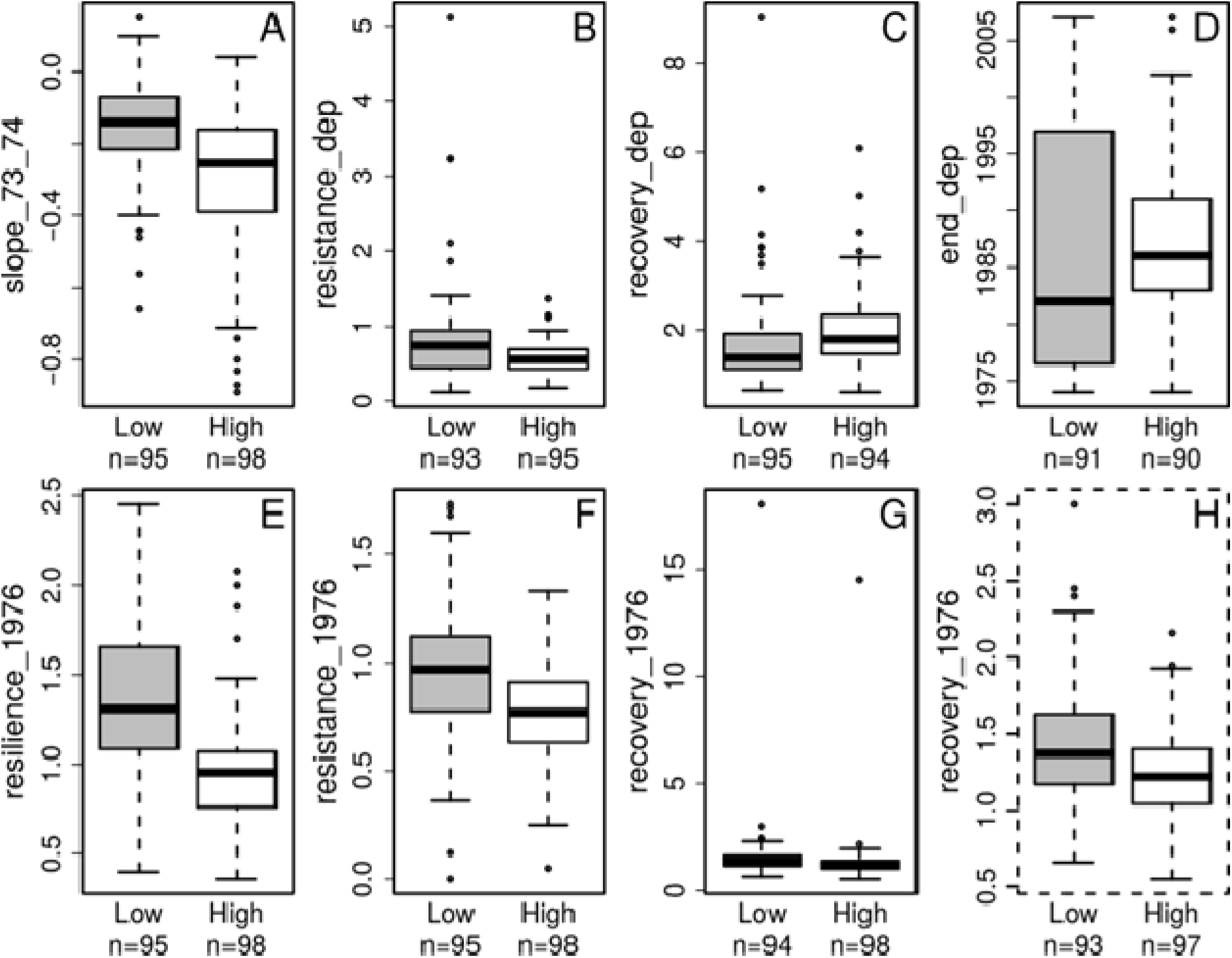
Dendrophenotypes for trees from both sites (Low and High). Phenotypes that describe the start (A) and end of the depression period (D), and the trees’ resistance (B) and recovery (C) are depicted. Resilience (E) and resistance (F) to and recovery (G) from the drought year 1976 are depicted. For a better visualization of the data distributions, the extreme values for recovery_1976 were removed (H).

### Population structure

Neither the STRUCTURE analysis, nor the PCA provided any indication for population substructure. The visual inspection of the bar plots in STRUCTURE clearly showed that almost all individuals were assigned to both clusters in a scenario with K = 2 without any apparent pattern (Fig. S1), and Ln P (D) declined steadily with increasing K (Fig. S2). In accordance, point clouds resulting from the PCA showed no apparent difference between the sampling sites (Fig. S3).

### Genetic association

The different association methods provided us with largely different numbers of SNPs associated to our dendrophenotypes (Table 2, Table S2). VSURF identified 10 to 22 SNPs for every dendrophenptypic trait with the exception of *recovery_1976*, for which VSURF only found one SNP. Boruta detected a lower number of SNPs, ranging from zero to six, which were also detected by VSURF in most cases. The GLM in TASSEL yielded no significant results (permutation p-value < 0.007).

**Table 2.**
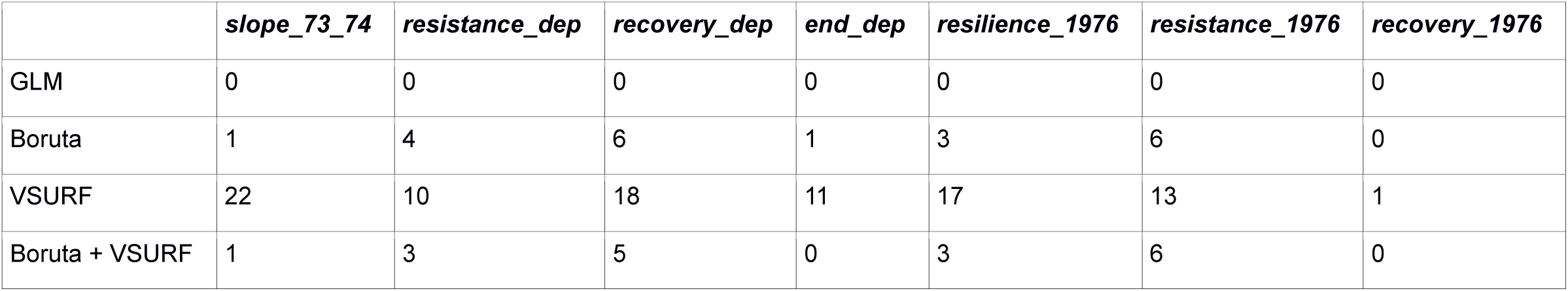
Overview of the results of the different association methods (Tassel GLM, Boruta and VSURF) for each dendrophenotype. Values in cells indicate the number of SNPs identified by each method for a given dendrophenotype.

In total, 15 SNPs were jointly identified by at least two approaches. Most of these SNPs are located in genes that code for membrane proteins related to transport and stress reactions (Table 3).

**Table 3.**
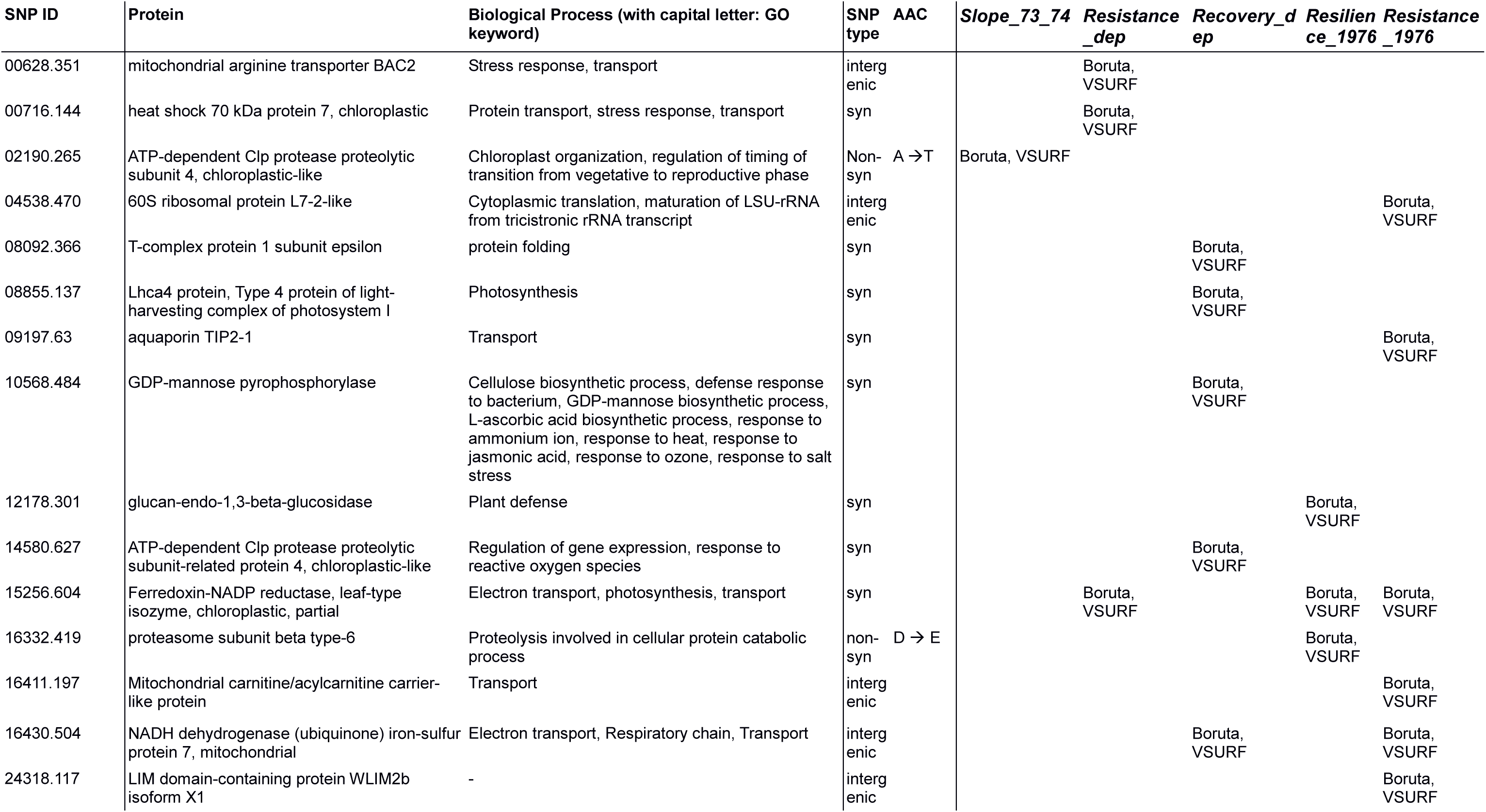
SNPs associated with scaled dendrophenotypes using Boruta and VSURF. Only SNPs that were associated with a given dendrophenotype with both methods are shown. A full table with all detected associations is provided in the supplementary material (Table S2). SNP IDs refer to the contigs and position of the SNP within the respective contig in the assembly of Roschanski et al. (2013). SNP type indicates whether a SNP is synonymous (syn), non-synonymous (non-syn), or located in intergenic regions. For nonsynonymous SNPs, the Amino acid change (AAC) is indicated.

| SNP ID | Protein | Biological Process (with capital letter: GO keyword) | SNP type | AAC | Slope_73_74 | Resistance_dep | Recovery_dep | Resilience_1976 | Resistance_1976 |
| --- | --- | --- | --- | --- | --- | --- | --- | --- | --- |
| 00628.351 | mitochondrial arginine transporter BAC2 | Stress response, transport | intergenic |  |  | Boruta, VSURF |  |  |  |
| 00716.144 | heat shock 70 kDa protein 7, chloroplastic | Protein transport, stress response, transport | syn |  |  | Boruta, VSURF |  |  |  |
| 02190.265 | ATP-dependent Clp protease proteolytic subunit 4, chloroplastic-like | Chloroplast organization, regulation of timing of transition from vegetative to reproductive phase | Non-syn | A → T | Boruta, VSURF |  |  |  |  |
| 04538.470 | 60S ribosomal protein L7-2-like | Cytoplasmic translation, maturation of LSU-rRNA from tricistronic rRNA transcript | intergenic |  |  |  |  |  | Boruta, VSURF |
| 08092.366 | T-complex protein 1 subunit epsilon | protein folding | syn |  |  |  | Boruta, VSURF |  |  |
| 08855.137 | Lhca4 protein, Type 4 protein of light-harvesting complex of photosystem I | Photosynthesis | syn |  |  |  | Boruta, VSURF |  |  |
| 09197.63 | aquaporin TIP2-1 | Transport | syn |  |  |  |  |  | Boruta, VSURF |
| 10568.484 | GDP-mannose pyrophosphorylase | Cellulose biosynthetic process, defense response to bacterium, GDP-mannose biosynthetic process, L-ascorbic acid biosynthetic process, response to ammonium ion, response to heat, response to jasmonic acid, response to ozone, response to salt stress | syn |  |  |  | Boruta, VSURF |  |  |
| 12178.301 | glucan-endo-1,3-beta-glucosidase | Plant defense | syn |  |  |  |  | Boruta, VSURF |  |
| 14580.627 | ATP-dependent Clp protease proteolytic subunit-related protein 4, chloroplastic-like | Regulation of gene expression, response to reactive oxygen species | syn |  |  |  | Boruta, VSURF |  |  |
| 15256.604 | Ferredoxin-NADP reductase, leaf-type isozyme, chloroplastic, partial | Electron transport, photosynthesis, transport | syn |  |  | Boruta, VSURF |  | Boruta, VSURF | Boruta, VSURF |
| 16332.419 | proteasome subunit beta type-6 | Proteolysis involved in cellular protein catabolic process | non-syn | D → E |  |  |  | Boruta, VSURF |  |
| 16411.197 | Mitochondrial carnitine/acylcarnitine carrier-like protein | Transport | intergenic |  |  |  |  |  | Boruta, VSURF |
| 16430.504 | NADH dehydrogenase (ubiquinone) iron-sulfur protein 7, mitochondrial | Electron transport, Respiratory chain, Transport | intergenic |  |  |  | Boruta, VSURF |  | Boruta, VSURF |
| 24318.117 | LIM domain-containing protein WLIM2b isoform X1 | - | intergenic |  |  |  |  |  | Boruta, VSURF |

## Discussion

In our study, the large majority of the investigated silver fir trees in the Bavaria Forest showed a pronounced growth decline from 1974 until the mid-1980s. Growth of most trees was reduced even further during the drought year in 1976. Although we did not find differences at the genetic level between our sampling sites, the growth decline was more severe at the high site as reflected in the dendrophenotypic traits measures. When we jointly analyzed the dendrophenotypic and genetic data, and simultaneously considered the effects of multiple SNPs with a random forest approach, we found that the variation in five out of seven dendrophenotypic traits was associated to the allelic variation of SNPs in 15 candidate genes.

### Dendrophenotypes

As expected, we found a pronounced population level growth decline in the 1970s and 1980s and population level recovery thereafter, as described earlier for silver fir in southern Germany (Elling *et al*., 2009). While our dataset is limited to surviving trees, inventory data from the area showed that silver fir dieback was substantial in the 1970s and 1980s. For example, some forest stands in the Bavarian Forest lost more than three-fourths of 80-120 year-old silver firs (unpublished inventory data, draft of the National Park Plan 1992). Although 1976 was not detected in an analysis for so-called pointer years (Cropper, 1979), we found that the 1976 drought had a negative effect on silver fir growth, particularly at the high site. As the methods to determine pointer years take a number of prior years as reference they are likely to fail if the previous years are already marked by reduced growth.

In contrast to 1976, dry years that did not coincide with the depression period (e.g. 1959, 1972, 1982 and 2003) did not have a strong effect on silver fir growth (Fig. 3). This is in line with Elling *et al*. (2009), who argued that SO_2_ pollution not only causes direct harm to silver fir trees by impeding photosynthesis and leading to the shedding of needles but that it also increases sensitivity to drought, which might be attributable to damages of the fine-root system.

In general, we observed that trees at high elevations were affected more severely during the depression period, as reflected by the generally lower resistance and resilience. However, without on-site measurements of SO_2_ concentrations, we can only speculate that the high site might have been more severely affected by SO_2_ emissions or, alternatively, that the effect was aggravated by site-specific conditions around the lake Rachel (e.g. higher air humidity).

As shown for other Central European silver fir stands, growth rates after the depression reached an unprecedented level, which is usually attributed to a combination of less dense forest structure after the dieback in the 1970s, as well as elevated nitrogen supply and increasing temperatures (Pinto *et al*., 2007; Elling *et al*., 2009; Büntgen *et al*., 2014). It has also been speculated that tropospheric ozone (O_3_) might be a major contributor to forest decline (Krause *et al*., 1986; Schmieden & Wild, 1995). However, since tropospheric O_3_ concentrations increased well into the 1980s, our data do not directly indicate a major influence on growth in silver fir stands in southern Germany.

### Dendrophenotype-genotype Association

So far, a few studies have jointly analysed genetic and dendroecological data in natural populations to explore the relationship between basic genetic parameters and growth traits (e.g. Pluess & Weber, 2012; King *et al*., 2013; Babushkina *et al*., 2016). None of the studies found a strong genetic signal related to the investigated growth traits, which could either be attributed to stronger effects of the environmental signals compared to the genetic influence on growth processes, or to a lack of adequate genetic data (e.g. loci that are relevant for the phenotypic traits considered). To take this one step further, we correlate dendrophenotypes and variation at stress response candidate genes (Roschanski *et al*., 2016) using a genetic association approach.

To our knowledge this is the first association study that links dendrophenotypes with SNPs in stress-related candidate genes. At our study sites, SNPs in 15 of the 103 candidate genes associated significantly with individual dendrophenotypic traits. Many of the genes were membrane proteins of the chloroplast, mitochondria or tonoplast, and thus, tightly linked to photosynthesis or chloroplast development. For example, SNPs in contigs 716 and 14580 which are associated to *resistance_dep* and *recovery_dep*, respectively, encode for stromal 70 kDa heat shock-related protein and a proteolytic subunit of the ATP-dependent Clp protease. Both are involved in protein folding with effects on chloroplast development and function (Sjögren *et al*., 2006; Latijnhouwers *et al*., 2010). Since SO_2_ pollution likely has negative effects on photosynthesis (Silvius *et al*., 1975), genes involved in these pathways could potentially determine how individual trees cope with these extreme conditions. Two of the genes that were exclusively associated with *resistance_1976* and *resilience_1976* can be directly related to drought stress response: aquaporin TIP2-1 and glucan-endo-1,3-beta-glucosidase. Aquaporins are regularly involved in drought response (Maurel *et al*., 2008; Hamanishi & Campbell, 2011) and a similar aquaporin (TIP1-1) has already been identified as differentially expressed in response to drought stress in silver fir seedlings (Behringer *et al*., 2015). Glucan-endo-1,3-beta-glucosidase was previously used as a drought stress candidate gene in *Pinus pinaster* (Eveno *et al*., 2008) and was also differentially expressed in response to drought stress in silver fir seedlings (Behringer *et al*., 2015). Two previous studies investigated local adaptation in silver fir along altitudinal gradients in France (Roschanski *et al*., 2016) and Southern Europe (Brousseau *et al*., 2016). Both studies identified a subset of the SNPs that showed patterns of divergent selection or correlated with environmental variables. The SNPs that associated with the dendrophenotypes did not overlap with the SNP that showed evidence of directional selection in France. Yet, two of the genes that were associated with *resistance_1976* and *resilience_1976* (contigs 04538 and 16332), were among the SNPs that were considered to be under divergent selection in the study of Brousseau et al. (2016) which provides additional evidence that these play a role in adaptation to extreme environmental conditions.

Eight of the SNPs found in associations are synonymous. Nevertheless, these mutations might impact gene expression, and it has been shown that there is a codon bias in conifers which affects translational efficiency (De La Torre *et al*., 2015).

### Statistical analyses of dendrophenotypes in association studies

Generally, dendrophenotypes are often complex phenotypes influenced by many genes with small effects. Classical single locus approaches will be less powerful to detect the underlying genetic signal. Therefore, it is key to utilize approaches that simultaneously consider the effects of multiple SNPs. Here, we applied a random forest based feature selection to identify SNPs that are likely associated with certain dendrophenotypes. The Boruta algorithm provides significant results by testing if the importance of a SNP for explaining a dendrophenotype is significantly (α = 5 %) higher than the importance of the most important randomly permuted attribute, which, under the null hypothesis is only associated by chance (Kursa *et al*., 2010). In contrast, VSURF does not incorporate any formal statistical hypothesis-test, but selects the most important SNPs regarding the association with a specific dendrophenotype (Genuer *et al*., 2015). However, this does not imply a statistically significant association. Both feature selection techniques are wrappers for the random forest algorithm and, as such, the importance value is a measure for the marginal effect of a SNP, as well as the interaction effect of all SNPs under consideration. It should be mentioned, however, that the relative contribution of marginal and interaction effect cannot be directly determined. Thus, SNPs identified by random forest procedures do not represent a network and have to be viewed independently. The benefit of such analyses is, however, that the influence of all other SNPs are incorporated in the importance of any given SNP, which provides a much better representation, given that in association studies of conifers, a single SNP never explained more than 5 % of the variation of a given trait (e.g. González-Martínez *et al*., 2006, 2008; Eckert *et al*., 2009).

### Outlook: The future of dendrophenotypes in association studies

Episodic environmental extremes like droughts, storms or other calamities are predicted to increase in both intensity and frequency due to global climate change (IPCC, 2014). A better understanding of the response of trees to such extreme events is urgently needed in order to predict their response to future climatic conditions. The unforeseeable timing of such extreme events makes it almost impossible to integrate them in research projects with short funding periods. However, as we have clearly shown in the present work, the time series nature of dendrophenotypes permits to characterize past tree responses to climate in general as well as to particular events, such as the response to extreme episodes. In addition, wood cores can be collected with an acceptable investment of time and money in the field and allow for diverse subsequent analyses of dendrophenotypic traits. These include anatomical features such as cell wall thickness or lumen area which are considered as proxy for physiological adaptations to external factors (Carrer *et al*., 2016; Ziaco *et al*., 2016) as well as isotope measures, which, for example, can be used to characterize the water use efficiency of a tree (Seibt *et al*., 2008). Microdensitometry can supplement ring width data with information of wood density, and thereby provide a more complete picture of growth for example during extreme events (e.g. Martinez-Meier *et al*., 2008). All these measures can be derived for long time series or with a focus on particular years of interest, providing exciting prospects for future studies. To facilitate this kind of research dendroecologists should start to focus on individual tree signals and not discard them as noise per default. Forest geneticists on the other hand should focus on pathways and candidate genes that are potentially related to dendrophenotypes.

## Contributions to Manuscript or Data

design of the research: BF, BZ, GV, KH, LO, SL, UB
performance of the research: CB, HJ, KH, LO, MW, SVL, UB
data analysis and interpretation: AP, DB, KH, LO, MW, SVL, UB
writing the manuscript: AP, ANP, BF, BZ, CB, DB, GV, KH, LO, PA, SVL, SL, UB

## Acknowledgements

We thank the Bavarian Forest National Park for supporting the field work. KH was funded by the ERAnet BiodivERsA project ‘TipTree’ (ANR-12-EBID-0003 granted to BZ, LO, BF, GV, SL) funded by the German Federal Ministry of Education and Research (Grant 01LC1202A to). MW and SVL were supported by ERASMUS scholarships during their stay at WSL Birmensdorf. UB received additional funding from the Ministry of Education, Youth and Sports of CR within the National Sustainability Program I (NPU I), grant number LO1415. Thanks to Lena Hellmann, Diego Galván Candela and Ricardo Ochoa Pereira for their help during field work, and to Anne Verstege for her support with tree-ring measurements.

